# Machine vision based frailty assessment for genetically diverse mice

**DOI:** 10.1101/2024.10.13.617922

**Authors:** Gautam S. Sabnis, Gary A. Churchill, Vivek Kumar

## Abstract

Frailty indexes (FIs) capture health status in humans and model organisms. To accelerate our understanding of biological aging and carry out scalable interventional studies, high-throughput approaches are necessary. We previously introduced a machine vision-based visual frailty index (vFI) that uses mouse behavior in the open field to assess frailty using C57BL/6J (B6J) data. Aging trajectories are highly genetic and are frequently modeled in genetically diverse animals. In order to extend the vFI to genetically diverse mouse populations, we collect frailty and behavior data on a large cohort of aged Diversity Outbred (DO) mice. Combined with previous data, this represents one of the largest video-based aging behavior datasets to date. Using these data, we build accurate predictive models of frailty, chronological age, and even the proportion of life lived. The extension of automated and objective frailty assessment tools to genetically diverse mice will enable better modeling of aging mechanisms and enable high-throughput interventional aging studies.

## 2 Introduction

Aging is a terminal process that is highly heterogeneous [1, 2]. Frailty is used to quantify this biological age and its heterogeneity. It is defined as the state of increased vulnerability to adverse health outcomes [3]. It is a clinically important measure in humans and is used preclinically in rodents in numerous aging studies. The frailty index (FI) is used to quantify frailty [4, 5]. An individual is scored on a set of age-related health deficits to produce a cumulative score [4]. Individual elements in a FI are selected to survey a broad set of organismal systems. With the exception of quantitative measures of body weight and core body temperature, all other characteristics were assessed on a scale of 0, 0.5, or 1 for normal, partly affected, and affected, respectively, by a trained technician blinded to genotype or age [2, 6]. All item scores are then summed to produce the cumulative FI score (CFI). Currently, FI scores outperform other measures, including molecular markers, in the prediction of mortality risk and health status [5, 7, 8].

The mouse frailty index represents a significant advancement in aging research, particularly for longterm interventional studies conducted across multiple laboratories. However, the FI scoring process requires trained personnel, hindering the tool’s scalability. Manually scoring FI for thousands of mice is labor-intensive, thus limiting the size of studies employing FIs [9]. Additionally, concerns regarding scorer-based variability and reproducibility arise due to the inherent subjectivity associated with several FI metrics [10-12]. While studies generally report good inter-scorer reliability for FI, they also highlight the crucial role of inter-scorer discussion and refinement in achieving high agreement [10-12]. This process may not be practical in multi-site or long-term studies. For instance, in our FI dataset of 515 mice, we found that the 42% of the phenotypic variation is use to the tester [13]. Furthermore, certain frailty items are more subjective than others. Therefore, while FI serves as a highly valuable tool for aging research, automating the scoring process to improve objectivity, scalability, reliability, and reproducibility would significantly enhance its overall utility.

In a previous paper, we used machine vision, the combination of computer vision and machine learning, to develop a video based automated frailty index, which we called a *visual frailty index* (vFI) [13]. Using video data from an open field assay, we extracted morphometric, gait, and other behavioral features that correlate with manual FI score and age. We use these features to train a regression model that accurately predicts the normalized FI score within 0.04 ±0.002 (mean absolute error). Unnormalized, this error is 1.08 ±0.05, which is comparable to 1 FI item being mis-scored by 1 point, or 2 FI items mis-scored by 0.5 points. The vFI provides increased reproducibility and scalability that will enable large-scale mechanistic and interventional studies of aging in mice. Overall, our work provided strong evidence that biological aging produces changes in behavior and physiology that are encoded in video data, i.e. we can visually determine the frailty of an animal based on their open field behavior. However, the vFI was constructed using data from 643 1-hr open field assays from 533 C57BL/6J (B6J) mice. Thus, the current vFI is constructed using only a single inbred mouse strain.

The importance of integrating genetic diversity in laboratory studies is widely recognized, including by the aging research community [14-17]. A B6J specific vFI is highly useful for mechanistic and interventional studies using this reference strain. However, understanding how genetic variation regulates complex aging phenotypes requires the ability to carry out frailty assessments in genetically diverse populations [18-20]. Indeed, studies of long-lived pedigrees indicate that 15-25% of the variation in lifespan is due to genetic factors [21-24]. Here, we extend the vFI to genetically diverse mouse populations. We collect manual FI data as well as open field video on 313 Diversity Outbred (DO) mice, and build models to predict chronologial and biological age, as well as proportion of life lived.

The DO is an advanced mouse population that originates from eight founder strains [25, 26]. There are approximately 45 million segregating single nucleotide polymorphisms (SNPs) in this population, much higher than classical laboratory strains [27, 28]. The population has been used successfully to model many complex phenotypes, including aging [29, 30]. DO mice have been used to study various aspects of aging including effect of diet, fasting, health span, life span, changes in home cage behavior, among others [31-37]. We hypothesize that mechanistic and interventional studies using genetically diverse mice are more likely to capture heterogeneity in aging due to genetics and generalize to humans. In addition to frailty, we are able to predict chronological age (vFRIGHT) and proportion of life lived (vPLL) using video data. The extension of video-based frailty assessment to genetically diverse mice greatly expands the utility of this noninvasive, objective, and sensitive tool.

## 3 Results

### 3.1 Exploratory Data Anlaysis

#### 3.1.1 Data collection and study design

Our workflow is shown in Figure 1A. Mice were tested in the open field and then assessed for manual frailty by experts. Using this data, we carried out a set of modeling experiments that are described in Figure 1B. The input to the models were either manual FI item scores or video features. From these inputs, we predicted chronological age (FRIGHT and vFRIGHT), biological age (vFI), or proportion of life lived (mPLL and vPLL) using statistical machine learning. Where possible, we attempted to compare performance of manual FI items with video features in the prediction task. We follow bestpractices for predictive statistical modeling and use multiple models, LM (penalized linear regression), random forest (RF), and XGBoost (XGB). We also report multiple performance metrics in the main text and supplementary figures MAE (median absolute error), RMSE (root mean square error), and R^2^ (Figures 4, 5, 6, S1). We describe each in detail below.

**Figure 1:**
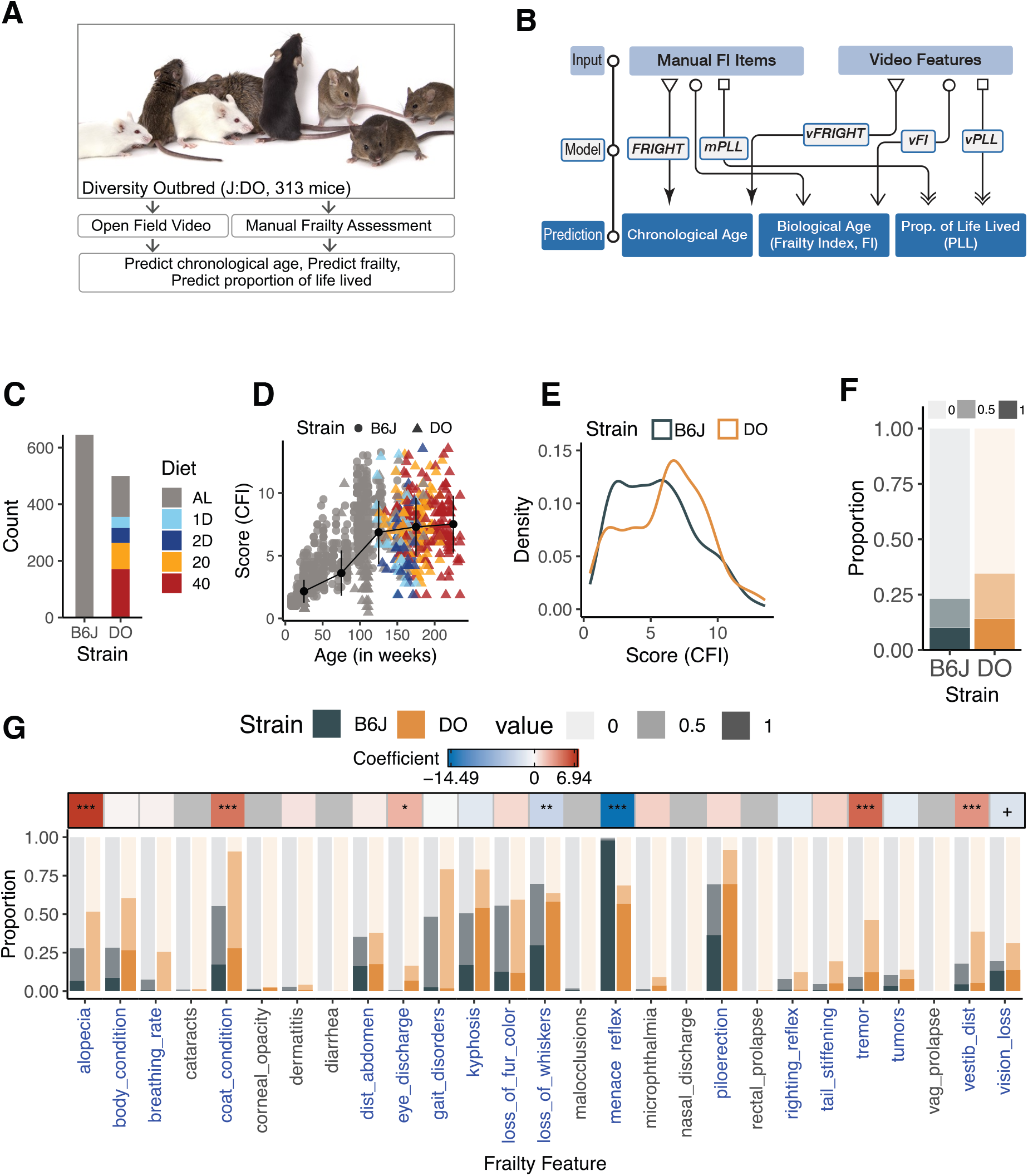
Overview of the study and exploratory frailty data analysis. (A) We use Diversity Outbred (DO) mice to predict chronological age, biological age, and predict proportion of life lived. We combine previous data from C57BL/6J mice that was published in [13] photo credit: JAX) (B) Description of our models. We have two sets of inputs, manual frailty items and video features. We describe 5 models that predict either chronological age, biological age, or percent life lived. (C) Summary of the number of animals per strain and diet. (D) Distribution of cumulative frailty index score (CFI) score by age. The black line shows a piece-wise linear fit (***n =*** 126,260,335,252,172 mice) to the data. The center point is the mean, and the error bars are the s.d. values. Color and shape indicate diet and mouse population, respectively (E) The cumulative frailty index score (CFI) or frailty index score (FI) distribution across B6J and DO populations. (F) Effect of scorer on FI data; 18% of the variability in manual FI scores was due to a scorer effect (RLRT = 9.22, ***p <*** 9 × 10^_4^). (G) Individual FI item usage in B6J and DO populations. We fit a cumulative link model for each frailty feature separately to examine the effect of strain on the manual scores after adjusting for weight and diet. The positive coefficients (red) indicate that a mouse belonging to the DO population increases the score, i.e., manual scores in the higher categories (0.5 or 1) are more likely, and vice versa for negative coefficients (blue). We used FDR-based correction of the Wald-based p-values for tests of the strain effect being zero for different frailty features. We colored a feature blue if at least 10% of individuals in either population show variation in manual score.

#### 3.1.2 Exploratory analysis of manual FI data

We analyze three populations: B6J, DO, and combined B6J-DO (ALL) datasets as shown in Table 1. The DO dataset was part of an ongoing study that has been previously described [31, 33, 36, 37]. We combined the DO dataset with a previously published B6J dataset for modeling and analysis [13]. The DO data comprises 500 data points collected on 313 unique DO mice. The DO mice were assigned to one of the five diets: ad libitum feeding (AL), fasting one day per week (1D) or two consecutive days per week (2D), and caloric restriction at 20% (20) or 40% (40) of baseline ad libitum food intake. The combined B6J-DO data contains 1145 data points collected longitudinally on 828 unique mice (515 B6J, 313 DO) (Figure 1C). DO population was highly imbalanced in favor of females. We recorded a top-down video of each mouse in a 1-hour open-field session according to previously published protocols [13, 38, 39]. Following the open-field recording, a trained expert scored and assigned a manual FI score. Consistent with previous data, in the B6J-DO dataset, we found that the mean FI score and the heterogeneity of the FI score increase with age (Figure 1D). The cumulative frailty index is largely identical between the two populations (Figure 1E).

**Table 1:** Summary table of animals and frailty scores in our dataset.

| Strain | Num. animals | Num. tests | M | F | Age | Frailty |
| --- | --- | --- | --- | --- | --- | --- |
| DO | 313 | 500 | 21 | 292 | 25 - 234 weeks | 0.5 - 14 |
| B6J | 515 | 645 | 299 | 216 | 8 - 161 weeks | 1 - 13.25 |

We explored the usage of individual manual frailty index (FI) items between B6J and DO populations. Given the genetic heterogeneity of DO, we hypothesized differential FI item usage. First, we examined the overall distribution of the scores (0, 0.5, 1) and observed no significant changes (Figure 1F) which is reasonable since the CFI distribution is similar between populations. We then looked at the usage of individual FI elements (Figure 1 G). An item is considered useful if at least 10% of individuals in either population show some variation (Figure 1G, blue vs. black). Some FI items are rarely used in both populations, items such as cataracts, dermatitis, diarrhea, and rectal or vaginal prolapse show inadequate variation. Overall, 18 of 27 (67%) of the FI items show sufficient variation to be considered useful (Figure 1G, Blue). We then identified which of these 18 have significantly different score distributions (0, 0.5, 1) between B6J and DO (Figure 1G, Top Ribbon). Seven of these 18 items show differential use, suggesting frailty manifests differently in the two populations. For example, coat condition, alopecia, and piloerection differ in frailty scores between B6J and DO.

### 3.2 Effect of environmental factors on manual frailty data

Next, we explored sources of variation in the manual FI data. A significant issue with manual FI scores is the variability introduced by different scorers. Previously, a scorer effect of 42% was observed in our B6J manual FI data [13] (reproduced in Figure S1A). We also detected a notable scorer effect in the DO FI data (18%, Figure 2A) and consequently in the combined B6J-DO FI data (26%, Figure 2A). For comparison, scorer-related variability in the manual FI data (18%) surpasses the variability due to diet (14%) in the DO data that includes severe calorie restriction. Since sex was highly imbalanced in favor of females (21M, 292F, Table 1), we did not analyze variation due to sex. To eliminate the tester effect from manual FI scores, we subtracted the scorer effects estimated using a linear mixed model (see Section 5) from the manual FI scores to obtain scorer-adjusted FI scores. In the following modeling sections, we constructed clocks for scorer-adjusted FI score prediction.

**Figure 2:**
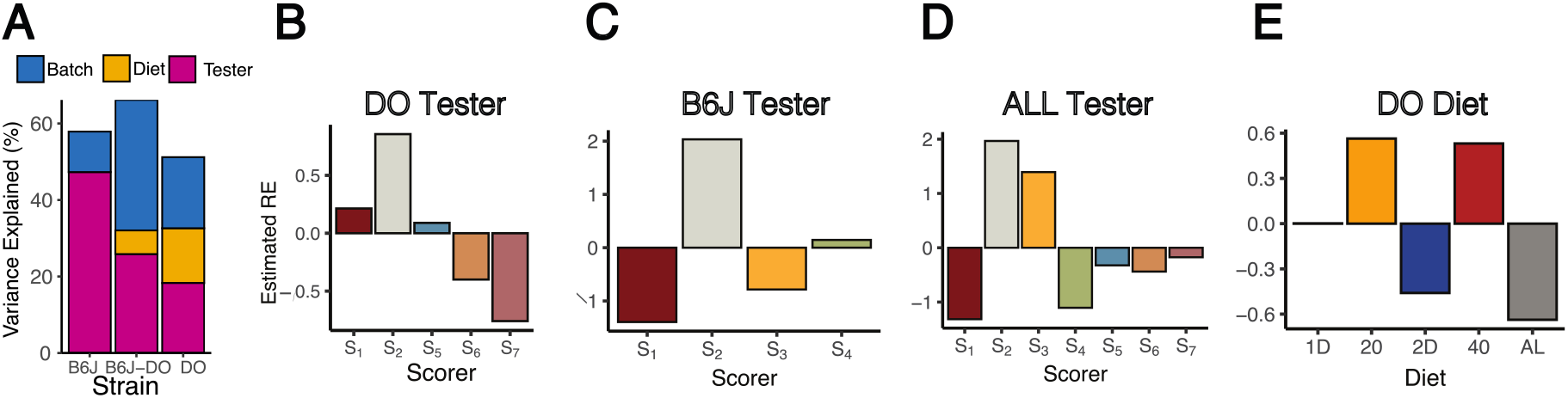
Variance due to tester, diet, and batch in the manual frailty data. (A) Using linear mixed models, we obtained the variability in scores (%) due to three factors (the residual variation (RE) is not plotted). We plotted the estimated random effects across scorers in (B) DO, (C) B6J, (D) B6J-DO, and across diets in the (D) DO dataset.

### 3.3 Open field video data processing and exploratory data analysis

We processed open-field videos using tracking and pose estimation networks to produce an ellipse fit and a 12-point mouse pose for each frame as previously described [13, 39]. Next, we used these frame-level measurements to calculate various per-video features, including traditional open-field measures, gait and postural measures, and engineered features using the JAX Animal Behavior System (JABS) [13, 39, 40]. Altogether, we extracted 59 features in three feature sets, open field, gait/posture, and engineered [13]. We determined whether features within and across three sets provide similar or independent information through a correlation matrix (Figure 3A). We found the presence of correlated blocks along the diagonal of the correlation matrix, indicating that measures within a feature set are more correlated than across the feature sets. We handled the presence of multicollinear features within and across sets through regularization of our predictive models in order to increase predictive accuracy.

**Figure 3:**
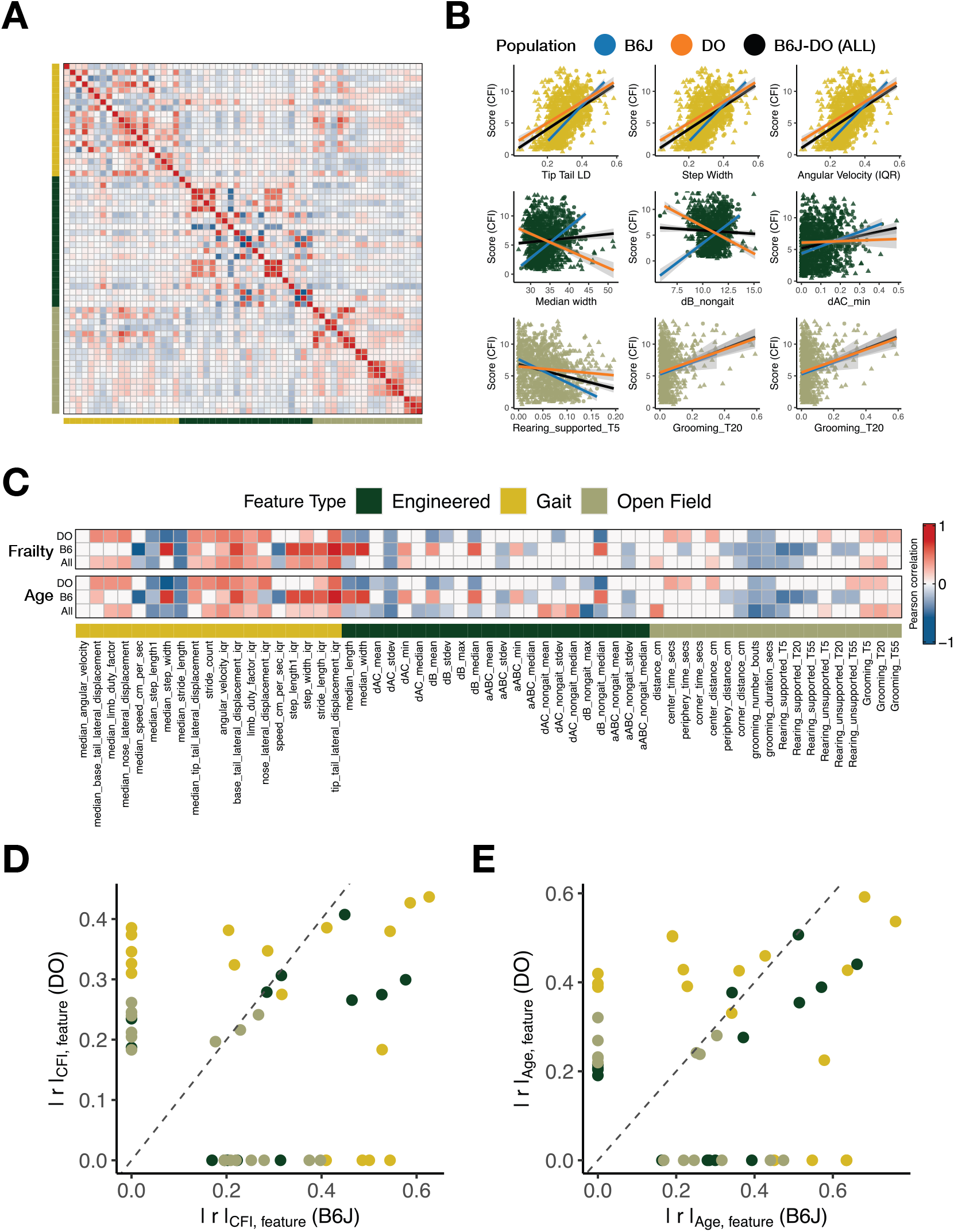
Exploratory analysis of the video features. (A) The estimated Pearson correlation matrix across all video features captures the linear interdependence (relationship) across multiple video features in our dataset. The diagonal elements show the variance of each individual variable, whereas the off-diagonal elements capture the correlation between each pair (row, column) of color-coded variables. (B) Each row corresponds to a feature type; the column corresponds to data B6J (left), DO (middle), and combined (right). We made a scatterplot for each feature type and strain combination and overlayed the best linear fits (B6J (blue), DO (orange), combined (black)) for the feature having the maximum correlation with the score (CFI). (C) Pearson correlations of all video features with frailty score (CFI) (top) and age (bottom). (D) A scatterplot showing features that we found to correlate with score (left) and age (right) specifically for B6J (x-axis), DO (y-axis), and combined datasets (dashed diagonal line).

Prior to modeling, we explored the relationships between the video features and the age and the manual frailty score (Cumulative Frailty Index, CFI). The goal of this analysis was to determine which video features correlate with age and frailty in the three populations. We calculated the feature correlations for all three populations, using frailty scores and age (Figure 3C). We find that overall, similar correlations are observed in the three goups. However, we also find distinct feature usage in the three populations. We plot a set of features that show similar correlations in all three populations (Figure 3B, top row), a set that show opposite correlation in B6 and DO populations (Figure 3B, middle row), and a set that shows correlations in a subset of three populations (Figure 3B, bottom row). To further explore this population-specific feature correlations, we plot the correlation of each video feature with CFI and age for the B6J and DO populations (Figure 3D,E, respectively). We find a group of video features that are exclusive to DO (y-axis aligned) and B6J (x-axis aligned). In addition, we find a set of commonly used features (middle of scatter plot). Surprisingly, we see population specific feature correlations across the three sets of video features (open field, gait, and engineered). These results led us to hypothesize that population specific models are needed to predict frailty, age, and proportion of life lived. Therefore, in all our anaysis we construct a B6J, DO, and B6J-DO (ALL) population specific models.

Using this data, we carried out a set of predictions that are described in Figure 1B. The input to the models were either manual FI item scores or video features. We predicted chronological age, biological age, or proportion of life lived using the FRIGHT, mPLL, vFRIGHT, vFI, or vPLL models. Where possible, we attempted to compare performance of manual FI items to video features in prediction task. We describe each in detail below.

### 3.4 Prediction of Chronological Age

We constructed clocks to predict chronological age (vFRIGHT clock) and biological age (vFI clock) from video features in the DO population. We labeled these clocks as DO-vFRIGHT and DO-vFI, respectively. We successfully constructed accurate clocks for the isogenic mouse strain B6J, namely B6J-vFRIGHT and B6J-vFI [13]. We showed that, in the B6J population, video features can accurately predict the animal’s age and frailty and outperform manual frailty items in predicting the animal’s age. We hypothesized that 1) B6J-specific clocks are not transferable to the DO population and would perform poorly, 2) DO-specific clocks trained on a genetically heterogeneous mouse population will be more accurate, and 3) a combined B6J-DO clock would perform better than individual population-specific clocks.

To test our first two hypotheses for chronological age prediction, we trained the B6J-vFRIGHT clock on theB6J data. Weaskedhow well the B6J trained vFRIGHT clock predicted the age of DO data [13]. The B6J-vFRIGHT clock predicted the DO mice’ age with an MAE of 75.8 weeks, an RMSE of 85.9 weeks, and *R*^*2*^ of 0.03 (Table S1). In contrast, the DO-vFRIGHT clock, trained and tested on DO mice using cross-validation, predicted the DO mice’ age with an MAE (median absolute error) of 14 ±0.06 weeks (Figure 4B), RMSE (root mean square error) of 18.7 ±0.05 weeks (Figure S1C), and *R*^2^ of 0.86 ±0.06 (Figure S1A). These results showed that the models explicitly trained on the DO population are far more accurate and that B6J-specific clocks are not transferable to the DO population. Next, we asked the reverse question of how well the DO-vFRIGHT clock can predict the age of B6J mice. We found it performed worse than the B6J-vFRIGHT clock, with an MAE of 74.6 weeks versus 12.1 weeks (B6J-FRIGHT clock) and similar poor performance in RMSE and *R*^2^ (Table S1). We plotted the predicted versus actual values for the held-out test folds for the B6J (Figure 4C) and DO (Figure 4D). We obtained similar results for the vFI clocks shown later (Section 3.5).

**Figure 4:**
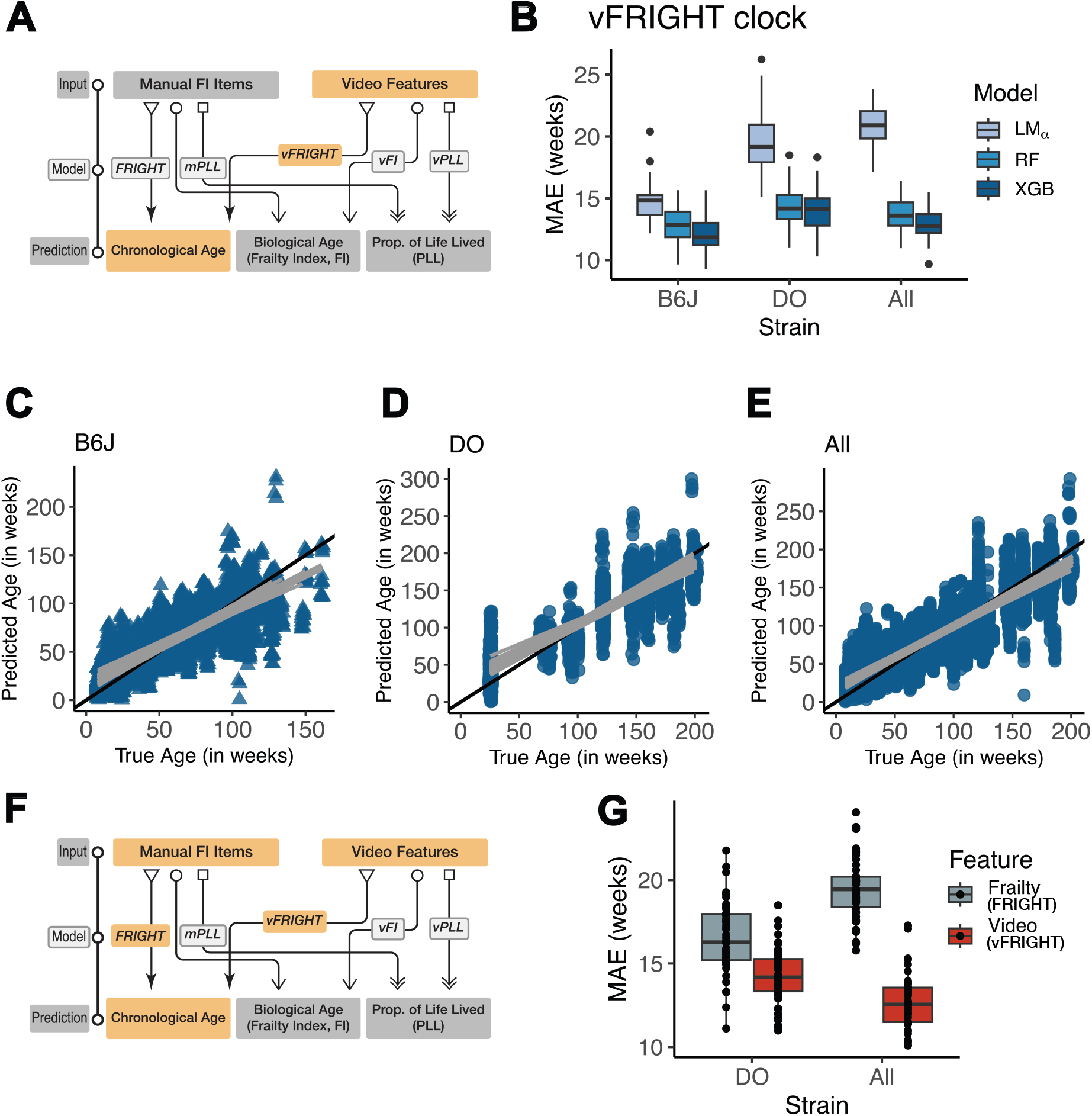
Age prediction using video features (vFRIGHT clock). (A) Modeling workflow for constructing vFRIGHT clock. (B) We tested three models (LM (penalized linear regression), random forest (RF), and XGBoost (XGB)) using 10 × 5 nested cross-validation (outer folds = 10, inner folds = 5). First, we tuned the model hyperparameters on the inner folds. Then, we performed model selection using the tuned models on the outer folds using mean absolute error (MAE, y-axis) across three datasets (x-axis) separately. (C) Scatterplots of predicted age versus true chronological age for B6J, (D) DO, and (E) combined from the best model (color-coded) obtained via model selection. Points represent individual folds from each of the 50 independent test sets. Lines represent linear regression for each of the 50 independent folds. (G) Modeling workflow for constructing FRIGHT clock. (H) Video features (vFRIGHT) outperform manual frailty features (FRIGHT) in predicting age. Each dot represents the error of one fold in a 50-fold cross validation setup.

To determine whether a combined B6J-DO model would outperform a single population-specific model, we combined the B6J-DO data to construct the best-performing B6J-DO vFRIGHT clock. We found that the B6J-DO clock performed similarly to the other two models, with an MAE of 13 ±1.3 weeks (Figure 4B), RMSE of 17.4 ±2.05 weeks (Figure S1C), and *R*^2^ of 0.89 ±0.03 (Figure S1A). We plotted the predicted versus actual values for the held-out test folds for the B6J-DO model(Figure 4E).

In addition to building the vFRIGHT clocks using the (v)ideo features for the individual and combined populations, we constructed FRIGHT clocks using manual frailty items as features (Figure 4F). Data from B6J mice showed that our vFRIGHT model could more accurately and precisely predict age than the FRIGHT clock. Our vFRIGHT versus FRIGHT comparisons revealed similar results in both DO data and the combined B6J-DO data. In the DO data, we found that the video features outperform manual frailty items in predicting age with an MAE of 14.2 ± 0.24 weeks versus 16.5 ± 0.3 weeks, RMSE of 18.7 ± 0.39 weeks versus 24.5 ± 0.52 weeks, and *R*^2^ of 0.89 ± 0.01 versus 0.76 ± 0.02 (Figure 4G). Similarly, for the combined B6J-DO data, we found that the video features outperformed manual frailty items in predicting age, i.e. MAE of 12.7 ± 0.23 versus 19.5 ± 0.26, RMSE of 17.6 ± 0.34 versus 27.5 ± 0.36 and *R*^2^ of 0.89 ± 0.004 versus 0.74 ±0.007 (Figure 4G).

### 3.5 Prediction of Frailty (Biological Age)

Next, we addressed the goal of constructing frailty clocks: predicting manual FI scores using video features for the DO and B6J-DO mouse populations (Figure 5A). First, we asked how the B6J-vFI clock performed on the DO mice. We predicted the frailty of DO mice using the B6-vFI clock trained on B6J mice and obtained an MAE of 3.31, RMSE of 3.87, and R2 of 6 × 10”^6^ (Table S1). The B6J-vFI trained and tested on B6J mice using cross-validation had an MAE of 1.19 ±0.12. In contrast, the DO-vFI clock trained and tested on DO mice using cross-validation had an MAE of 1.53 ±0.18 (Figure 5B). These results validated our hypotheses that the B6J-vFI clock is not transferable and that a DO-vFI clock is more accurate in predicting the frailty of a DO mouse. The combined B6J-DO vFI clock had an MAE of 1.38 ±0.11 (Figure 5B), RMSE of 1.83 ± 0.52(Figure S1D), *R*^*2*^ of 0.70 ± 0.05(Figure S1B). The error, adjusted for the number of frailty items, is 0.06 (FI scores range can range from 0 to 1; our DO dataset had a range of 0.05 to 0.53). Compared to the manual FI items, an error of 1.6 is akin to either one FI item incorrectly scored as one and another FI item incorrectly scored as 0.5 and three items incorrectly scored at 0.5. We plotted the residuals for B6J, DO, and combined B6J-DO data to look at the predictions at the animal level and didn’t notice any irregularities (Figure 5 C,D,E).

**Figure 5:**
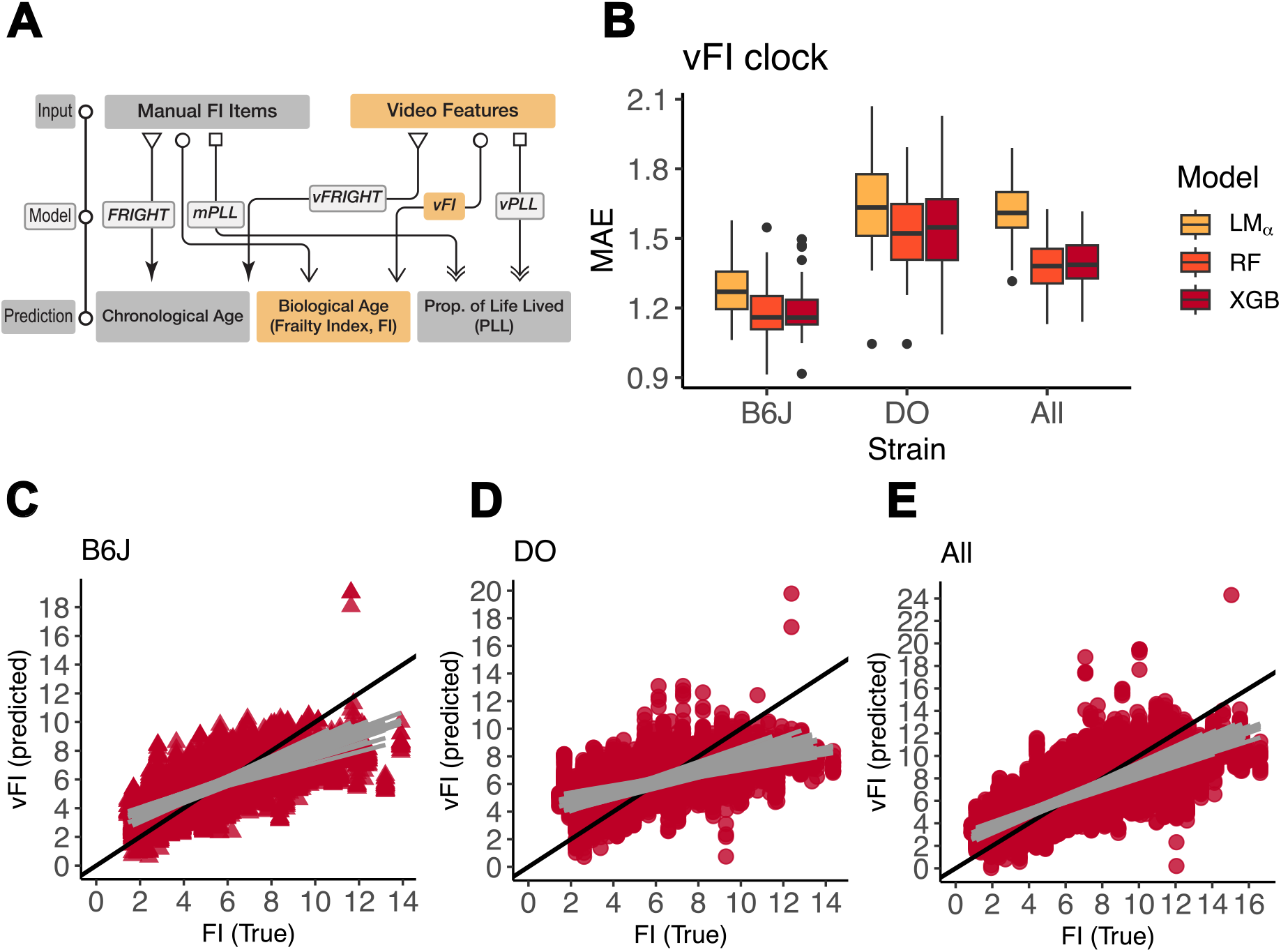
Biological age (frailty) prediction using video features (vFI clock). (A) Modeling workflow highlighting (in yellow) the clocks we construct using the video features as input to predict biological age (frailty). (B) We tested three models (LM (penalized linear regression), random forest (RF), and XGBoost (XGB)) using 10×5 nested cross-validation (outer folds = 10, inner folds = 5). First, we tuned the model hyperparameters on the inner folds. Then, we performed model selection using the tuned models on the outer folds using mean absolute error (MAE, y-axis) across three datasets (x-axis) separately. (C) Scatterplots of predicted age versus true chronological age for B6J, (D) DO, and (E) combined from the best model (color-coded) obtained via model selection. Points represent individual folds from each of the 50 independent test sets. Lines represent linear regression for each of the 50 independent folds.

### 3.6 Prediction of Proportion of Life Lived

Since our clocks can accurately predict age and frailty, we asked if we could also predict the animal’s mortality (Figure 6A). To answer this question, we defined the proportion of life lived (PLL) for each animal as the ratio of age at the test (in weeks) to age at death (in weeks). We used video and manual frailty items to compare their predictive performance in predicting PLL (Figure 6A). We used video features to predict PLL with an MAE of 0.072 ± 0.02, RMSE of 0.086 ± 0.25, and *R*^2^ of 0.37 ± 0.27. Similarly, we predicted PLL using manual frailty items with an MAE of 0.074 ± 0.020, RMSE of 0.09 ± 0.02, and *R*^*2*^ of 0.35 ± 0.25 (Figure 6B). We didn’t see an improvement in the performance using the combined set of features (video + frailty).

**Figure 6:**
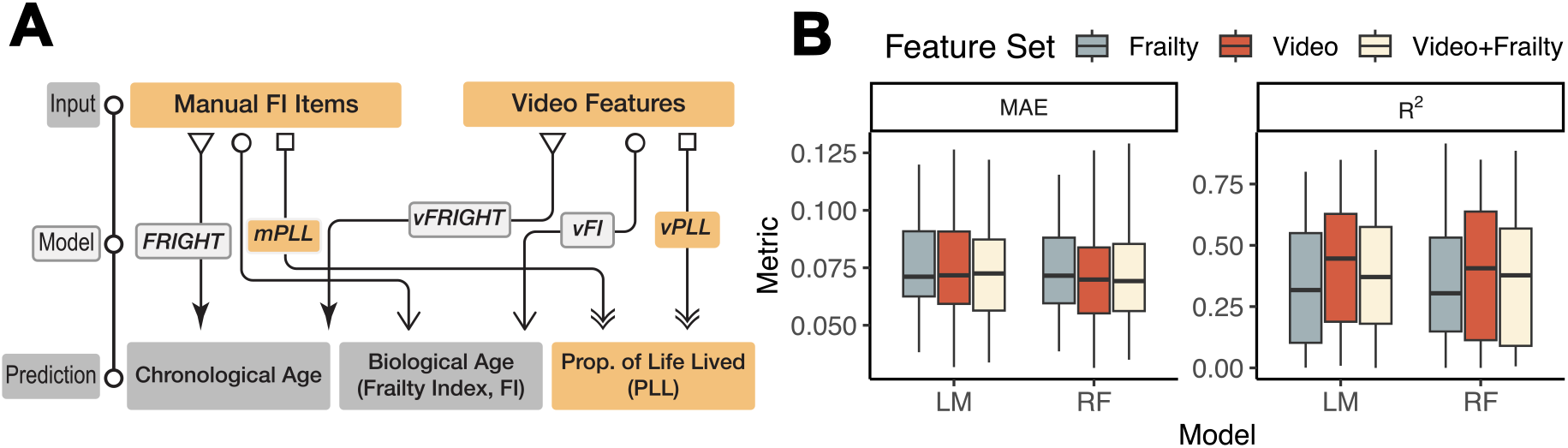
Proportion of Life Lived Prediction from manual frailty and video features. (A) Modeling workflow, highlighting (in yellow) the clocks we construct using the video and manual frailty features as input. (B) We tested two beta regression models; linear model (LM) and random forest (RF). The boxplots show the MAE and ***R***^**2**^ across of the held out folds in 50 fold cross-validation.

## 4 Discussion

The frailty index is an invaluable tool for aging research [9, 41, 42]. As the method becomes widely adopted, large centers such as the Nathan Shock Center for Aging Research at JAX and the Interventional Testing Program (ITP) will use it for longitudinal studies [15, 43]. These centers employ many trained staff for frailty assessments and often work on multiple multiyear studies simultaneously. Manual frailty assessment has subjectivity, even with trained staff, as testers can differ from each other and themselves over time. Recent work has shown that the professional background of scorers affects FI scores. Specifically, inter-scorer reproducibility was poor between animal technicians and research scientists [11]. These studies on inter-scorer agreement strongly emphasize the importance of interscorer discussion and refinement in obtaining high agreement [10-12]. In fact, it has been shown that in the absence of discussion and refinement, inter-rater reliability does not improve solely by practice and experience [11]. This discussion and refinement is not always feasible in multi-site or long-term studies. This variance can represent a critical challenge for aging studies in rodents that span several years. Indeed, in our previous B6J dataset consisting of 643 tests collected at the JAX Shock Center, we found that the tester accounted for 42% of the variability in manual FI scores. In the current DO dataset, 18% of the variation is due to the tester. For comparison, the variation due to diet, which introduces severe caloric restriction, was only 14%. Thus, the technical effect is larger than the diet effect.

We created the visual frailty assay (vFI) to provide an objective, sensitive, and scalable assay that can complement the manual frailty assay. We were quite surprised by how much information about chronological and biological age is encoded in the behavior of the animal. The vFI could accurately predict the frailty status of with an error of 1.08 ±0.05, which is comparable to 1 FI item being mis-scored by 1 point, or 2 FI items mis-scored by 0.5 points on our assay [13]. Our previous results also demonstrated that video was much more accurate in predicting chronological age than manual frailty items. One limitation of our work was that it used only the B6J strain of mice. Here, we extend our methods to genetically diverse mice.

Aging is highly genetic, and studies of long-lived individuals indicate that 15-25% of the variation in lifespan is due to genetic factors [21-24]. Studying aging in one inbred mouse strain, even if it is the reference strain, is akin to deeply studying one individual. Findings may not generalize to an entire population or across species. Impactful animal studies must incorporate genetic diversity into its design [18-20]. The DO and its companion Collaborative Cross (CC) are advanced mouse populations designed to model high genetic variance with balanced alleles. They are ideally suited for high-resolution genetic mapping and for modeling diverse populations for complex traits [25, 30]. We took advantage of an ongoing study at JAX to extend our behavior-based models to predict chronological age, biological age, and proportion of life lived. However, studying genetically diverse mouse populations introduces their own set of concerns, particularly for methods that use computer vision [38]. DO mice are highly genetically diverse, with concomitant phenotypic variation. This includes visual variation in size, coat color, as well as variation in behavior. In exploratory analysis, we observed population-specific usage of manual frailty items and video features. When we applied the previously trained B6J vFRIGHT and vFI model to the DO dataset, we saw poor performance. We concluded that population-specific models are needed for the prediction of chronological age, frailty, and PLL prediction.

Our models can accurately predict chronological age (vFRIGHT), frailty (vFI), and proportion of life lived (vPLL). The vFRIGHT and vFI clocks for DO have slightly worse performance than the B6J clocks, although the decrease in performance is incremental. We also compared the performance of FRIGHT and vFRIGHT clocks to compare the predictive value of manual frailty items and video features, respectively, in chronological age prediction. Our previous analysis using B6J showed that the video features are more predictive of chronological age than the manual frailty items in B6J. This is also seen in DO models. PLL clocks, also called the FRIGHT clock [5], attempt to predict the normalized life expectancy of an animal. We find that both manual FI and vFI have a similar predictability of PLL (mPLL and vPLL).

Overall, we have now generated one of the largest datasets of frailty data in the mouse that includes data from inbred and outbred mice. The accompanying video data can be used to model with both supervised and unsupervised methods to derive new behavior-based features. Our vFI is constructed using open field data and as proof of principle that video encodes informative aging features. Although the vFI is an important advance, the democratization of this technology remains a barrier. The method requires behavioral testing expertise and equipment, as well as computational expertise to implement, i.e. it is not plug-and-play. Although these resources are generally available in behavioral neuroscience laboratories, many geroscientists lack access, which prohibits the adoption of the method. Future developments need to alleviate the need for computational and behavioral expertise. One possible solution is continual monitoring of frailty using digital cages. Video based home cage monitoring is a rapidly advancing field and can be adopted for aging studies [32, 44], although no commercial systems are available. Ideally, a home cage monitoring system can provide a continuous frailty assessment without the need for computational expertise throughout the life of the animal. Data from social behaviors, sleep/wake states, feeding/drinking, and nesting behaviors can be combined with gait, posture, and other information described for the vFI to derive this continuous scale. An advanced home-cage monitoring system would offload the data management and compute requirements for scalable analysis to cloud compute resources, since such platforms would create terabytes of video data. In fact, there is still a large room for innovation and improvement in these digital approaches.

## 5 Methods

All animals and methods have been previously described. We carried out behavior tests according to published methods [38-40, 45]. The DO diet intervention dataset has been previously described [33, 36, 37]. All animals protocols followed JAX IACUC guidelines and no new animals were generated for this study.

We estimated the tester effect from the FI scores using a linear mixed model with the lme4 R package [46]. We had five testers in our DO data, and each tester tested a different number of animals. We subtracted the tester effects, estimated with the best linear unbiased predictors using restricted maximum likelihood estimates, from the FI scores of the animals. We labeled them as the tester-adjusted FI scores, 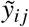.

We modeled the tester-adjusted FI scores with video-generated features using a linear regression model with elastic net penalty [47], random forest [48], and gradient-boosting machine [49] with trees as base learners. We trained several supervised machine learning models (Figure 4A) to test our hypotheses and constructed different sets of clocks. We tested three models: penalized linear regression (LR), random forest (RF), and extreme gradient boosting (XGB). We used a k-fold nested cross-validation resampling strategy for every model to tune model hyperparameters, perform model selection, and assess our models’ prediction accuracy and generalization performance to new unseen data. We ensured that the repeat measurements from the same animal belonged to either the training or the test data and not both. We used three performance measures to quantify how well the model predictions match the ground truth: mean absolute error (MAE), root-mean-squared-error (RMSE), and *R*^*2*^.

## 6 Acknowledgments

We thank J. Graham Ruby and Andrea Di Francesco, Calico Life Sciences LLC for sharing DO frailty data and advice on this manuscript. We thank Kumar Lab member Marina Santos for open field testing. We thank Shock Center and Churchill Lab members Hannah Donato, Gaven Garland, Mackenzie Leland, and Laura Robinson for frailty indexing and coordinating. We thank members of the Kumar Lab for critical feedback, including Brian Geuther, Jaycee Choi, Anshul Choudhary, and Jacob Beierle. We thank the members of the JAX Information Technology team for infrastructure support.

## 9 Supplement

**Figure S1:**
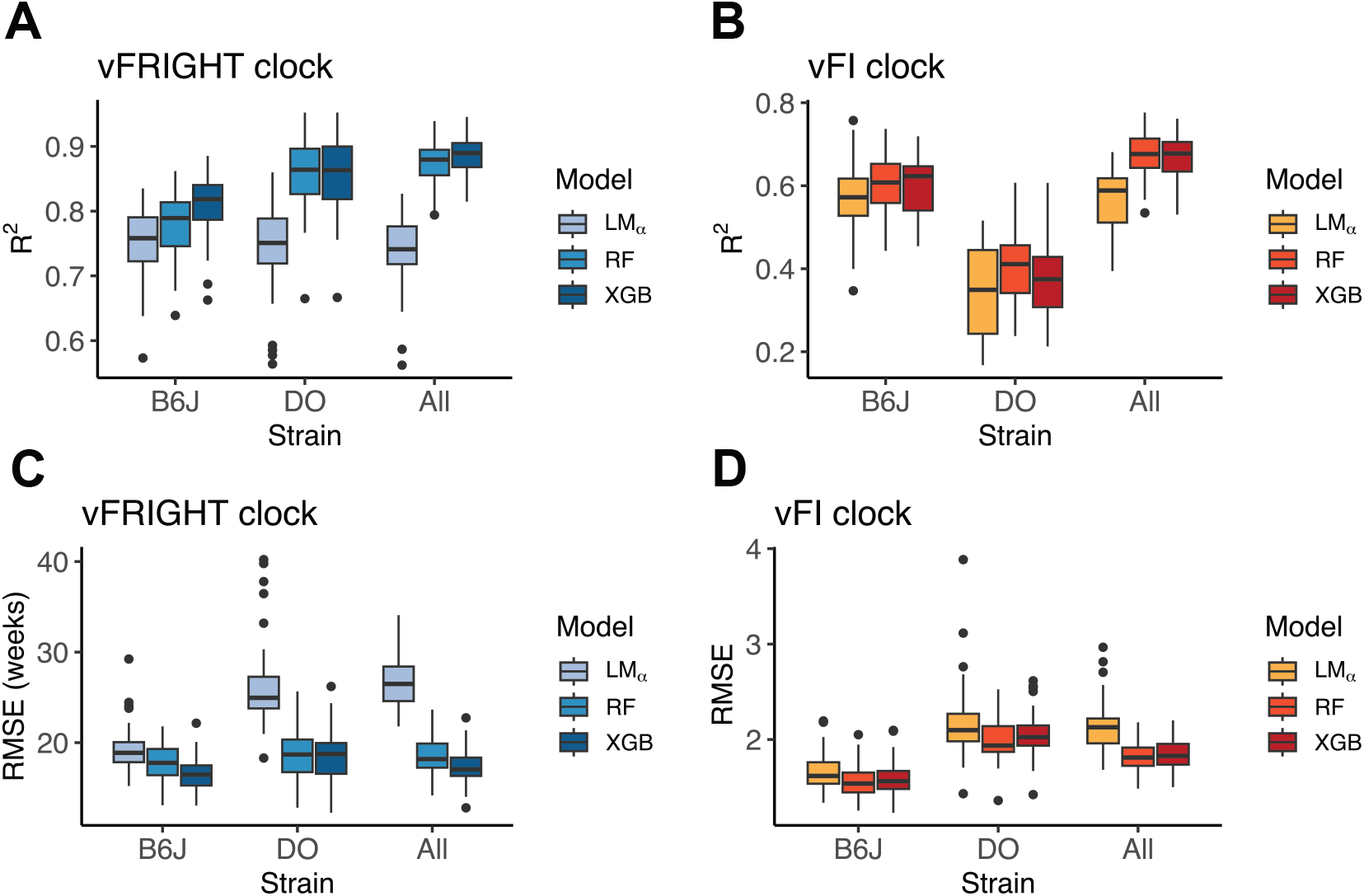
Modeling performance of vFRIGHT (A, C) and vFI clocks (B, D) quantified with R^2^ (A, B) and RMSE (C D) across different models and datasets.

**Table S1:** Cross performance of model trained on one population on the other. We trained vFRIGHT and vFI clocks (models) on the B6J and DO populations and tested their predictive performance on DO and B6, respectively. We found that the clocks did not generalize well to the mouse populations outside the training set, as evidenced by large mean absolute error (MAE), root mean squared error (RMSE), and high *R*^2^ values.

| Training | Inference | Model | MAE | RMSE | $R^2$ |
| --- | --- | --- | --- | --- | --- |
| B6J | DO | vFRIGHT | 75.8 (weeks) | 85.9 (weeks) | 0.03 |
| B6J | DO | vFI | 3.31 (unnormalized FI) | 3.87 (unnormalized FI) | $\approx 0$ |
| DO | B6J | vFRIGHT | 63.3 (weeks) | 74.6 (weeks) | 0.05 |
| DO | B6J | vFI | 2.26 (unnormalized FI) | 2.76 (unnormalized FI) | 0.06 |

## References

1. Mitnitski, A., Mogilner, A. & Rockwood, K. Accumulation of Deficits as a Proxy Measure of Aging. TheScientificWorldJournal 1, 323–36 (Sept. 2001).

2. Whitehead, J. C. et al. A clinical frailty index in aging mice: comparisons with frailty index data in humans. Journals of Gerontology Series A: Biomedical Sciences and Medical Sciences 69, 621–632 (2014).

3. Rockwood, K., Fox, R. A., Stolee, P., Robertson, D. & Beattie, B. L. Frailty in elderly people: an evolving concept. CMAJ 150, 489–495. ISSN: 0820-3946 (1994).

4. Searle, S. D., Mitnitski, A., Gahbauer, E. A., Gill, T. M. & Rockwood, K. A standard procedure for creating a frailty index. BMC geriatrics 8, 24 (2008).

5. Schultz, M. B. et al. Age and life expectancy clocks based on machine learning analysis of mouse frailty. Nature communications 11, 1–12 (2020).

6. Sukoff Rizzo, S. J. et al. Assessing healthspan and lifespan measures in aging mice: optimization of testing protocols, replicability, and rater reliability. Current protocols in mouse biology 8, e45 (2018).

7. Kim, S., Myers, L., Wyckoff, J., Cherry, K. E. & Jazwinski, S. M. The frailty index outperforms DNA methylation age and its derivatives as an indicator of biological age. English. GeroScience 39, 83–92 (Jan. 2017).

8. Kojima, G., Iliffe, S. & Walters, K. Frailty index as a predictor of mortality: a systematic review and meta-analysis. Age and Ageing 47, 193–200. ISSN: 0002-0729 (Oct. 2017).

9. Rockwood, K. et al. A frailty index based on deficit accumulation quantifies mortality risk in humans and in mice. Scientific reports 7, 43068 (2017).

10. Feridooni, H. A., Sun, M. H., Rockwood, K. & Howlett, S. E. Reliability of a Frailty Index Based on the Clinical Assessment of Health Deficits in Male C57BL/6J Mice. The Journals of Gerontology: Series A 70, 686–693. ISSN: 1079-5006 (Sept. 2014).

11. Kane, A. E. et al. Factors that Impact on Interrater Reliability of the Mouse Clinical Frailty Index. The Journals of Gerontology: Series A 70, 694–695. ISSN: 1079-5006 (Apr. 2015).

12. Kane, A. E., Ayaz, O., Ghimire, A., Feridooni, H. A. & Howlett, S. E. Implementation of the mouse frailty index. Canadian journal of physiology and pharmacology 95, 1149–1155 (2017).

13. Hession, L. E., Sabnis, G. S., Churchill, G. A. & Kumar, V. A machine-vision-based frailty index for mice. Nature aging 2, 756–766 (2022).

14. Flurkey, K., Currer, J. M. & Harrison, D. in The mouse in biomedical research 637–672 (Elsevier, 2007).

15. Korstanje, R., Peters, L. L., Robinson, L. L., Krasinski, S. D. & Churchill, G. A. The Jackson Laboratory Nathan Shock Center: impact of genetic diversity on aging. Geroscience 43, 2129– 2137 (2021).

16. Phelan, J. P. Genetic variability and rodent models of human aging. Experimental gerontology 27, 147–159 (1992).

17. Vanhooren, V. & Libert, C. The mouse as a model organism in aging research: usefulness, pitfalls and possibilities. Ageing research reviews 12, 8–21 (2013).

18. Curtsinger, J. W. et al. Genetic variation and aging. Annual review of genetics 29, 553–575 (1995).

19. Greenwood, P. & Parasuraman, R. Normal genetic variation, cognition, and aging. Behavioral and Cognitive Neuroscience Reviews 2, 278–306 (2003).

20. Melzer, D., Hurst, A. J. & Frayling, T. Genetic variation and human aging: progress and prospects. The Journals of Gerontology Series A: Biological Sciences and Medical Sciences 62, 301–307 (2007).

21. Newman, A. B. et al. A meta-analysis of four genome-wide association studies of survival to age 90 years or older: the Cohorts for Heart and Aging Research in Genomic Epidemiology Consortium. Journals of Gerontology Series A: Biomedical Sciences and Medical Sciences 65, 478–487 (2010).

22. Murabito, J. M., Yuan, R. & Lunetta, K. L. The search for longevity and healthy aging genes: insights from epidemiological studies and samples of long-lived individuals. Journals of Gerontology Series A: Biomedical Sciences and Medical Sciences 67, 470–479 (2012).

23. Schoenmaker, M. et al. Evidence of genetic enrichment for exceptional survival using a family approach: the Leiden Longevity Study. European Journal of Human Genetics 14, 79–84 (2006).

24. Broer, L. et al. GWAS of longevity in CHARGE consortium confirms APOE and FOXO3 candidacy. Journals of Gerontology Series A: Biomedical Sciences and Medical Sciences 70, 110–118 (2015).

25. Svenson, K. L. et al. High-resolution genetic mapping using the Mouse Diversity outbred population. Genetics 190, 437–447 (2012).

26. Churchill, G. A., Gatti, D. M., Munger, S. C. & Svenson, K. L. The diversity outbred mouse population. Mammalian genome 23, 713–718 (2012).

27. Yang, H. et al. Subspecific origin and haplotype diversity in the laboratory mouse. Nature genetics 43, 648–655 (2011).

28. Aylor, D. L. et al. Genetic analysis of complex traits in the emerging Collaborative Cross. Genome research 21, 1213–1222 (2011).

29. Schmidt, C. W. Diversity outbred: a new generation of mouse model 2015.

30. Saul, M. C., Philip, V. M., Reinholdt, L. G. & Chesler, E. J. High-diversity mouse populations for complex traits. Trends in Genetics 35, 501–514 (2019).

31. Di Francesco, A. et al. Dietary restriction impacts health and lifespan of genetically diverse mice. Nature. ISSN: 1476-4687. 10.1038/s41586-024-08026-3 (Oct. 9, 2024).

32. Ruby, J. G. et al. An Automated, Home-Cage, Video Monitoring-based Mouse Frailty Index Detects Age-associated Morbidity in C57BL/6 and Diversity Outbred Mice. The Journals of Gerontology: Series A 78, 762–770 (2023).

33. Wright, K. M. et al. Age and diet shape the genetic architecture of body weight in diversity outbred mice. Elife 11, e64329 (2022).

34. Di Francesco, A. et al. Regulators of health and lifespan extension in genetically diverse mice on dietary restriction. bioRxiv, 2023–11 (2023).

35. Hackett, S. R. et al. The Molecular Architecture of Variable Lifespan in Diversity Outbred Mice. bioRxiv, 2023–10 (2023).

36. Luciano, A. et al. Longitudinal fragility phenotyping predicts lifespan and age-associated morbidity in C57BL/6 and diversity outbred mice. Biorxiv (2024).

37. Prateek, G. et al. Longitudinal analysis of body weight reveals homeostatic and adaptive traits linked to lifespan in diversity outbred mice. bioRxiv, 2024–06 (2024).

38. Geuther, B. et al. Robust mouse tracking in complex environments using neural networks. Communications Biology 2, 124 (Mar. 2019).

39. Beane, G. et al. Video based phenotyping platform for the laboratory mouse. bioRxiv (2022).

40. Sheppard, K. et al. Stride-level analysis of mouse open field behavior using deep-learning-based pose estimation. Cell Reports 38, 110231 (Jan. 2022).

41. Banga, S., Heinze-Milne, S. D. & Howlett, S. E. Rodent models of frailty and their application in preclinical research. Mechanisms of ageing and development 179, 1–10 (2019).

42. Mishra, M. & Howlett, S. E. in Frailty: A Multidisciplinary Approach to Assessment, Management, and Prevention 81–89 (Springer, 2024).

43. Nadon, N. L., Strong, R., Miller, R. A. & Harrison, D. E. NIA interventions testing program: investigating putative aging intervention agents in a genetically heterogeneous mouse model. EBioMedicine 21, 3–4 (2017).

44. Baran, S. W. et al. Digital biomarkers enable automated, longitudinal monitoring in a mouse model of aging. The Journals of Gerontology: Series A 76, 1206–1213 (2021).

45. Kumar, V. et al. Second-generation high-throughput forward genetic screen in mice to isolate subtle behavioral mutants. Proceedings of the National Academy of Sciences 108, 15557–15564. ISSN: 0027-8424 (2011).

46. Bates, D., Maechler, M. & Bolker, B. Walker., S. Fitting linear mixed-effects models using lme4. J Stat Softw 67, 1–48 (2015).

47. Zou, H. & Hastie, T. Regularization and variable selection via the elastic net. Journal of the royal statistical society: series B (statistical methodology) 67, 301–320 (2005).

48. Breiman, L. Random forests. Machine learning 45, 5–32 (2001).

49. Friedman, J. H. Greedy function approximation: a gradient boosting machine. Annals of statistics, 1189–1232 (2001).

